# A phylogenomic framework for Vetigastropoda (Mollusca)

**DOI:** 10.1101/736447

**Authors:** Tauana Junqueira Cunha, Gonzalo Giribet

**Affiliations:** Harvard University

## Abstract

Abalones, turban snails, top snails, keyhole limpets and slit shells are just some of the diverse marine Vetigastropoda. With major lineages having ancient divergences in the Paleozoic Era, basal nodes in the phylogeny have been largely unresolved. Here we present the first genomic-scale dataset focused on vetigastropods, including a comprehensive sampling of taxa with all superfamilies and about half of the extant families (41 new transcriptomes, 49 total ingroup terminals). Our recovered topology is congruent among inference methods for the majority of nodes, and differs from previous studies, in which long branch attraction artifacts seem to be pervasive. The first three divergences sequentially separate Pleurotomarioidea, Seguenzioidea and Haliotoidea from remaining vetigastropods. Fissurelloidea is sister group to Scissurelloidea, Lepetodriloidea and Lepetelloidea, which all together are the sister group to Trochoidea. A methodological discordance exists in the position of Haliotidae and Fissurellidae, possibly due to a compounded effect of small sample bias derived from gene-wide partitioning. Dense sampling of trochoids and fissurellids also provides a backbone for family and subfamily-level relationships, with Tegulidae not monophyletic.

## 1.1 Introduction

Abalones, turban snails, top snails, keyhole limpets, slit shells and little slit shells are just some of the diverse marine Vetigastropoda. With about 3,700 living species, and many more thousands of fossil taxa, vetigastropods are one of the five major lineages of gastropods (Cunha and Giribet 2019), and the dominant gastropod clade throughout the Paleozoic and most of the Mesozoic eras (Fryda et al. 2008). They are distributed at all depths and types of habitats, from shallow hard substrates, to sand, to deep sea vents and cold seeps. Despite many attempts, an understanding of the evolutionary relationships between the main clades of vetigastropods remains a challenge. With such an old divergence of the major lineages, many morphological traits show convergence, and the amount of molecular data collected so far has not been sufficient to resolve short internodes in the tree.

The following eight superfamilies are currently accepted containing 38 extant families (Bouchet et al. 2017; WoRMS 2019): Pleurotomarioidea (Pleurotomariidae: slit shells), Fissurelloidea (Fissurellidae: keyhole and slit limpets), Haliotoidea (Haliotidae: abalones), Lepetelloidea (eight families: deep sea limpets), Lepetodriloidea (two families: hydrothermal vent limpets), Scissurelloidea (four families: little slit shells), Seguenzioidea (eight families), Trochoidea (13 families: turban and top snails).

Since Vetigastropoda was first recognized in 1980 (Salvini-Plawen 1980), there has been no shortage of phylogenetic studies on the group. Early on, comprehensive morphological analyses tried to establish relationships and identify synapomorphies (Haszprunar 1988; Ponder and Lindberg 1997; Salvini-Plawen and Haszprunar 1987; Sasaki 1998). On the molecular side, early studies used a handful of markers and provided incremental contributions to solving vetigastropod relationships (Geiger and Thacker 2005; Harasewych et al. 1997; Yoon and Kim 2005). Some key studies, though with few markers, presented densely sampled phylogenies, helping place not only the most diverse groups, but also minute and hard-to-collect taxa (Aktipis and Giribet 2011; Kano 2008; Williams et al. 2008). Finally, a recent burst of publications based on mitochondrial genomes have targeted deep vetigastropod relationships, albeit with limited taxon sampling (Lee et al. 2016; Uribe et al. 2016, 2017; Wort et al. 2017).

These efforts led to few stable results. Pleurotomariidae has been recovered as the sister group to all other vetigastropods since the first molecular phylogenies (Harasewych et al. 1997). This contradicted morphological hypotheses in which pleurotomariids were more closely related to other families with symmetric, paired pallial organs (e.g. haliotids, fissurellids, scissurellids) and to the asymmetric trochids (Ponder and Lindberg 1997; Salvini-Plawen 1980; Sasaki 1998). Trochoidea, the most diverse superfamily of vetigastropods, has also received considerable attention, with many dedicated studies (Uribe et al. 2017; Williams 2012; Williams et al. 2010, 2008; Williams and Ozawa 2006). Phasianellidae and Angariidae, once considered subfamilies of Turbinidae, have later been elevated to family and even superfamily status, being recovered as early divergences in the tree (Williams et al. 2008; Williams and Ozawa 2006). More recently, however, they have been consistently recovered within the superfamily Trochoidea with mitochondrial genomes and a small sample of vetigastropod transcriptomes (Lee et al. 2016; Uribe et al. 2016, 2017; Zapata et al. 2014).

Except for the position of Pleurotomariidae, and despite the numerous studies on Vetigastropoda and Trochoidea, relationships between superfamilies and families remain contentious. Most datasets have non-overlapping taxa representation and do not have enough sequence data to resolve such old divergences. In addition, mitochondrial genomes suffer from having fast evolutionary rates, which regularly create artifacts of long branch attraction (LBA) (Uribe et al. 2016, 2019). This issue is accentuated by insufficient taxon sampling of vetigastropods and outgroups. Particularly problematic taxa that have been placed in very different positions depending on the study, and that seem to be more affected by LBA, are Fissurellidae, Haliotidae, Lepetelloidea and Lepetodriloidea (Aktipis and Giribet 2011; Lee et al. 2016; Uribe et al. 2016, 2017; Williams et al. 2008). Familial relationships within Trochoidea have been redefined recently with relatively good support, but pending the inclusion of several missing families (Uribe et al. 2017).

Here we present new transcriptomes for 41 genera of vetigastropods, and combine them with previously published sequences. Our sampling covers all superfamilies and about half of the described families to resolve deep relationships at those levels. This is the first genomic-scale dataset focused on vetigastropods, including a large representation of taxa. Our recovered topology is well supported and congruent among inference methods for the majority of nodes.

## 1.2 Methods

### 1.2.1 Sampling and sequencing

We sequenced the transcriptomes of 41 vetigastropod genera and added another eight genera with previously published sequences, for a total of 49 ingroup terminals. We further used 20 outgroup gastropods and one bivalve outgroup, for a total of 62 terminals. This sampling covers all eight superfamilies of vetigastropods, and 18 of the 38 families. All new data and selected published sequences are paired-end Illumina reads. New samples were fixed in RNA*later* (Invitrogen). RNA extraction and mRNA isolation were done with the TRIzol Reagent and Dynabeads (Invitrogen). Libraries were prepared with the PrepX RNA-Seq Library kit using the Apollo 324 System (Wafergen). Quality control of mRNA and cDNA was done with a 2100 Bioanalyzer, a 4200 TapeStation (Agilent) and the Kapa Library Quantification kit (Kapa Biosystems). Samples were pooled in equimolar amounts and sequenced in the Illumina HiSeq 2500 platform (paired end, 150 bp) at the Bauer Core Facility at the Harvard University.

### 1.2.2 Transcriptome assembly and matrix construction

Raw reads were assembled *de novo* with Trinity v2.3.2 (Grabherr et al. 2011; Haas et al. 2013) with the same pipeline and scripts from Cunha and Giribet 2019. The completeness of the assemblies was evaluated with BUSCO v3.0.2 by comparison with the Metazoa database (Simão et al. 2015). Orthology assignment of the peptide assemblies was done with OMA v2.2.0 (Altenhoff et al. 2018). We built four matrices to account for extreme evolutionary rates, amino acid composition heterogeneity and two levels of matrix completeness. We first used a custom python script (*selectslice.py*) to select all orthogroups for which at least half of the terminals were represented (50% taxon occupancy), resulting in a matrix with 1027 genes (matrix 1) (Figure 1.1). Each orthogroup was aligned with MAFFT v7.309 (Katoh and Standley 2013), and the ends of the alignment were trimmed to remove positions with more than 80% missing data with a custom bash script (*trimEnds.sh*). A subset of 259 genes with 70% taxon occupancy constitutes matrix 2. To avoid possible biases, saturation and long-branch attraction, matrix 3 was built by removing from matrix 1 the 20% slowest and the 20% fastest evolving genes, as calculated with TrimAl (Capella-Gutierrez et al. 2009), for a final size of 615 genes (Figure 1.1). Matrix 4 is the subset of 894 genes from matrix 1 that are homogeneous regarding amino acid composition. Homogeneity for each gene was determined with a simulation-based test from the python package p4 (Foster 2004, 2018), with a custom script modified from Laumer et al. (2018) (*p4_compo_test.py*) and a conservative p-value of 0.1. For inference methods that require concatenation, genes were concatenated using Phyutility (Smith and Dunn 2008). We further reduced compositional heterogeneity in matrices 1 and 2 by recoding amino acids into the six Dayhoff categories (Dayhoff et al. 1978) with a custom bash script (*recdayhoff.sh*). All scripts are available from Cunha and Giribet 2019.

**FIGURE 1.1:**
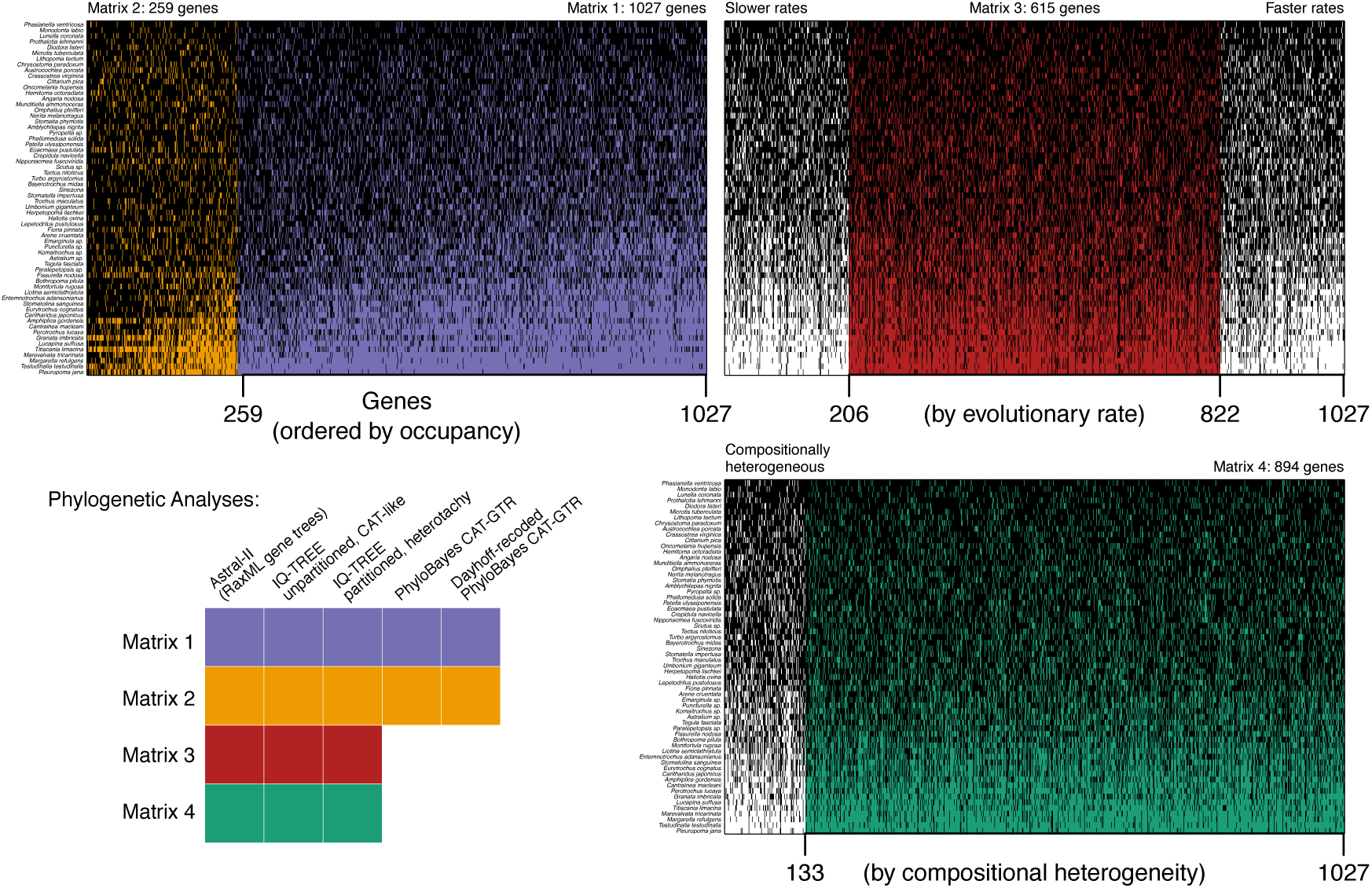
Matrices and methods used to infer vetigastropod relationships. Matrix 1 includes 1027 genes with 50% taxon occupancy. Matrix 2 is the subset of 259 genes with 70% taxon occupancy. Genes and species are sorted with the best sampling on the upper left. Matrix 3 is the subset of 615 genes after ordering all genes by evolutionary rate and removing the 20% slowest and 20% fastest evolving genes. Matrix 4 includes the 894 genes that are homogeneous in amino acid composition. Black cells indicate genes present for each species. Check Methods for details.

### 1.2.3 Phylogenetic analyses

Amino acid matrices were used for phylogenetic inference with a coalescent-based approach in Astral-II v4.10.12 (Mirarab and Warnow 2015), with maximum likelihood (ML) in IQ-TREE MPI v1.5.5 and v1.6.8 (Chernomor et al. 2016; Kalyaanamoorthy et al. 2017; Nguyen et al. 2015), and with Bayesian inference (BI) in PhyloBayes MPI v1.7a (Lartillot et al. 2013). The two Dayhoff-recoded matrices were analyzed in PhyloBayes (Figure 1.1). For the coalescent-based method, gene trees were inferred with RAxML v8.2.10 (Stamatakis 2014) (-N 10 -m PROTGAMMALGF) and then used as input for Astral-II for species tree estimation. For each concatenated matrix, we inferred the best ML tree with two strategies: a partitioned analysis with model search including LG4 mixture models and accounting for heterotachy (-bb 1500 -sp partition_file -m MFP+MERGE -rcluster 10 -madd LG4M,LG4X -mrate G,R,E) (the best-fit partitioning scheme maintained all genes as separate partitions); and a non-partitioned analysis with model search including the LG and WAG rate matrices with a profile mixture model (Le et al. 2008) (ML variant of the Bayesian CAT model (Lartillot and Philippe 2004)) (-bb 1500 -m MFP -mset LG,WAG -rcluster 10 -mfreq F+C20 -mrate G,R). Computer memory limited which of the profile models (C10 to C60) could be used depending on the size of the matrix. The following best-fit models were selected for each matrix: WAG+F+C20+R10 (matrices 1 and 4), WAG+F+C60+R7 (matrix 2), WAG+F+C30+R7 (matrix 3). PhyloBayes was run with the CAT-GTR model on matrices 1 and 2, discarding constant sites to speed up computation. Tree figures were edited with the R package *ggtree* (Yu et al. 2017).

## 1.3 Results

### 1.3.1 Phylogenetic analyses

Our 16 analyses included amino acid and Dayhoff-recoded matrices, four schemes of gene subsampling, and Bayesian inference, maximum likelihood and coalescent-based approaches to infer deep relationships between superfamilies and families of vetigastropods (Figure 1.1). The result is an overall well resolved and well supported phylogeny that covers all of the eight superfamilies and about half of the extant family diversity (Figure 1.2). Vetigastropoda is monophyletic, and Pleurotomarioidea is sister group to all other vetigastropods. Seguenzioidea, here represented by the family Chilodontaidae, then diverges from remaining groups, followed by the divergence of Haliotoidea (Figure 1.2). The remaining diversity is then split in two large clades, Trochoidea on one side, and Lepetelloidea, Lepetodriloidea, Scissurelloidea and Fissurelloidea as its sister group. Within the latter, Lepetelloidea (represented by Pseudococculinidae and Pyropeltidae) shares a more recent common ancestor with Lepetodriloidea (represented by Lepetodrillidae), and this clade is sister group to Scissurelloidea (represented by Scissurellidae) (Figure 1.2).

**FIGURE 1.2:**
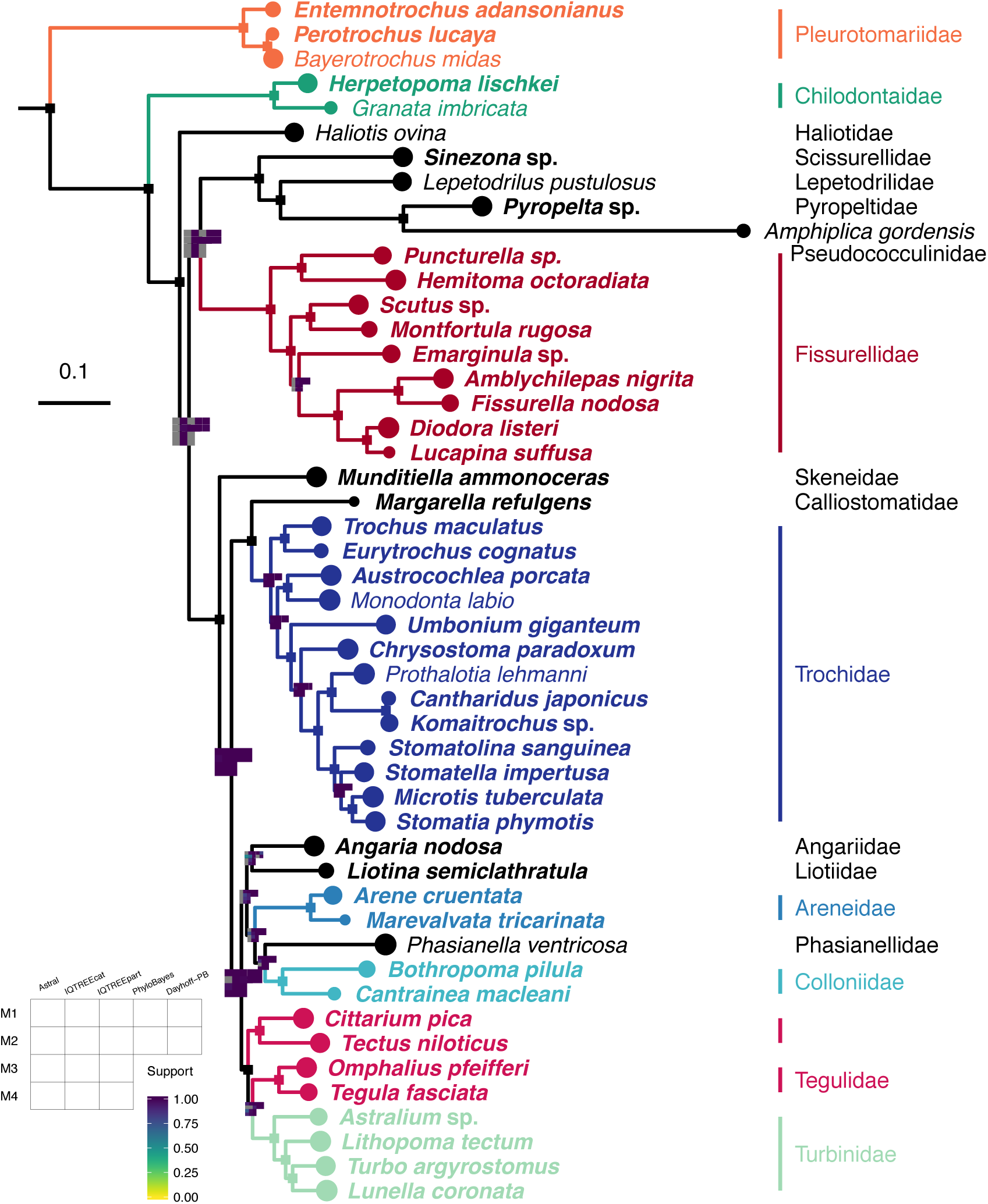
Vetigastropod phylogeny from matrix 1 with IQTREEcat: ML profile mixture model. Single squares mark branches with full support in all analyses; remaining nodes have plots colored according to support value (0-1) for each analysis. Grey squares represent splits that were absent in a given analysis. New transcriptomes in bold. M1-M4: matrices 1-4; IQTREEpart: ML partitioned; Dayhoff-PB: BI Dayhoff-recoded matrix. See Methods for details.

Within Trochoidea, Skeneidae is sister group to all other trochoid families. One clade then unites Calliostomatidae and Trochidae, and another encompasses the rest of the familial diversity of Trochoidea. Colloniidae and Phasianellidae are sister group to Areneidae, and this clade is recovered as the sister group of Angariidae and Liotiidae (Figure 1.2). Tegulidae is not monophyletic, with the clade containing *Tegula* being more closely related to Turbinidae than to other tegulids. With the exception of Tegulidae, all families with more than one sampled genus are monophyletic and well supported in all analyses. In Pleurotomariidae, *Perotrochus* and *Bayerotrochus* are more closely related than either is to *Entemnotrochus*. In Fissurellidae, Diodorinae and Fissurellinae are sister subfamilies, but Emarginulinae is not monophyletic, with *Emarginula* being more closely related to Diodorinae and Fissurellinae than to *Scutus* and *Montfortula*. This entire clade is then sister group to the subfamily Zeidorinae (*Puncturella* and *Hemitoma*) (Figure 1.2). In Trochidae, all subfamilies with more than one genus are monophyletic. Trochinae is the sister group to all other trochids, with Monodontinae then diverging from the remaining subfamilies. Stomatellinae (*Stomatella* and relatives) is the sister group to Cantharidinae (*Cantharidus* and relatives). These two subfamilies are sister to Chrysostomatinae (*Chrysostoma*), and the clade is sister group to Umboniinae (*Umbonium*) (Figure 1.2). In Turbinidae, *Astralium* is recovered as the sister group to other sampled genera, with *Lithopoma* then diverging from *Lunella* and *Turbo*.

Inference methods are largely congruent across the tree, however two nodes show conflicting results: the branch leading to the clade that is sister group to Haliotidae, and the branch leading to the clade of Scissurelloidea, Lepetodriloidea, Lepetelloidea and Fissurelloidea. In both cases, Bayesian inferences and maximum likelihood analyses with the more complex profile mixture model result in the tree shown in Figure 1.2 with full support (Figures A.1, A.2). On the other hand, coalescent-based analyses and most gene-partitioned maximum likelihood analyses result in a different topology, where Haliotoidea is the sister group to Trochoidea, this clade is sister group to Fissurellidae, and finally this large clade is sister group to the clade of Scissurelloidea, Lepetelloidea and Lepetodriloidea (Figures A.3, A.4).

## 1.4 Discussion

We present the first comprehensive phylogenomic framework for Vetigastropoda, including all super-families and about half of the family diversity (Figure 1.2). As expected based on previous molecular studies (Harasewych et al. 1997; Kano 2008; Williams et al. 2008), Pleurotomarioidea is the sister group to all other vetigastropods. Apart from that, previous work on this major gastropod lineage has resulted in many conflicting and poorly resolved topologies. Our resulting backbone of the vetigastropod tree is largely concordant among matrices and inference methods, but a methodological discordance still exists in two key basal nodes discussed below.

### 1.4.1 Methodological discordance

Short branches lead to the two basal nodes with conflicting results, which likely indicates this was a time of rapid divergence between vetigastropod clades, making it hard to infer the true history of divergences. Results did not differ based on the strategy of gene subsampling, but based on the model of inference that was used. Bayesian analyses, which used the CAT model for site heterogeneity, and the maximum likelihood analyses that similarly used a CAT-like profile mixture model both resulted in the tree shown in Figure 1.2. A different placement of Haliotoidea and Fissurelloidea is however recovered in analyses ran with gene-partitioned maximum likelihood and with the shortcut coalescent-based approach (Figures A.3, A.4; see section 1.3).

The discordance is interesting because some recent phylogenomic studies have found agreement between coalescent-based approaches and concatenated datasets analyzed with the more complex profile mixture models of sequence heterogeneity (Cunha and Giribet 2019; Schwentner et al. 2017). Approaches based on the multispecies coalescent account for incomplete lineage sorting and therefore are theoretically more sound (Degnan and Rosenberg 2009; Edwards 2009). The agreement with some concatenated datasets indicates that more complex models were effectively accounting for the heterogeneity present in such large concatenated matrices. In practice, however, fully coalescent approaches that infer gene trees and the species tree simultaneously are computationally intensive and currently impossible to run on densely-sampled genomic datasets. Shortcut methods were then developed that use maximum likelihood gene trees to infer the species tree, accelerating computation time (Mirarab and Warnow 2015). One of the issues of shortcut methods is therefore gene tree estimation error, derived from systematic inference error or from insufficient information contained in short alignments (Meiklejohn et al. 2016; Molloy and Warnow 2018; Sayyari et al. 2017).

It has recently been shown that excessive partitioning leads to a compounded effect of small sample bias over many single protein partitions (Wang et al. 2019). This in turn results in a marked long branch attraction (LBA) issues for analyses using protein-wise partitioning (Wang et al. 2019). Because shortcut coalescent approaches such as Astral rely on single-gene analyses, they are also expected to be susceptible to errors based on gene partitioning. We argue that the congruence between our partitioned ML and coalescent results indicates a likely bias of gene-wise partitioning and LBA. Interestingly, it has been observed that using gene trees with less informative alignments caused a shift towards more pectinate (asymmetric) species trees (Meiklejohn et al. 2016). This was also observed in our results, with the alternative position of Fissurelloidea (Figures A.3, A.4), producing a more asymmetric tree than the one shown in Figure 1.2. Modeling site-heterogeneity is one of the most important strategies to avoid biases in large genomic datasets (Meiklejohn et al. 2016; Schwentner et al. 2018), and it has recently been recommended that gene-partitioning should be avoided due to strong LBA biases (Wang et al. 2019). We therefore consider the results of our site-heterogeneous models to provide a more accurate representation of the true evolutionary history of vetigastropods at the conflicting nodes (Figure 1.2).

### 1.4.2 Relationships among vetigastropod superfamilies

We find a well supported and stable clade composed of Scissurelloidea, Lepetodriloidea and Lepetelloidea (Figure 1.2). Because of the minute size of scissurelloids and the deep sea environments inhabited by the two latter superfamilies, these taxa have been some of the hardest to sample in molecular phylogenies (but see Kano 2008). Lepetelloidea in particular is absent in most studies, but has been recovered as the sister group to Patellogastropoda, which was nested in Vetigastropoda, in a seven-gene phylogeny (Aktipis and Giribet 2011). The unexpected position of both clades and their exceptional long branches in that tree indicate a clear artifact of LBA. Here we find that Lepetelloidea is closely related to Lepetodriloidea and to other superfamilies in the order Lepetellida, in agreement with current classification (Bouchet et al. 2017; WoRMS 2019). Scissurelloidea and Lepetodriloidea have been recovered as sister groups in many molecular studies in which Lepetelloidea was absent (Aktipis and Giribet 2011; Geiger and Thacker 2005; Kano 2008; Williams et al. 2008; Williams and Ozawa 2006; Yoon and Kim 2005), which is concordant with our results. In mitochondrial genome analyses, Scissurelloidea has not been sampled yet, and Lepetodriloidea is in turn placed with Haliotoidea and Seguenzioidea (Lee et al. 2016; Uribe et al. 2016). These three superfamilies display very long branches in mitogenome studies (Lee et al. 2016; Uribe et al. 2016), again indicating a likely effect of LBA.

Seguenzioidea (here represented by Chilodontaidae) has been recovered with Haliotoidea in several mitogenome and nuclear studies (Aktipis and Giribet 2011; Lee et al. 2016; Uribe et al. 2016, 2017) and more broadly with Haliotoidea, Scissurelloidea and Lepetodriloidea in other phylogenies (Kano 2008; Williams et al. 2008). Here this superfamily is the sister group to all vetigastropods except pleurotomariids (Figure 1.2). Our results could be interpreted as an artifact of LBA, pushing Seguenzioidea to diverge earlier in the tree. However, its branch is not particularly long, and the position of this superfamily is completely stable and fully supported across all our analyses, many of which use site-heterogeneous models that are known to be more robust against LBA artifacts (Lartillot et al. 2007; Le et al. 2008). Interestingly, Seguenziidae has been placed in a similar position in early morphological studies (Ponder and Lindberg 1997; Sasaki 1998). This group has had various placements even more broadly in the gastropod phylogeny, sometimes being considered as more closely related to Caenogastropoda due to similarly complex reproductive anatomy (reviewed in Kano 2008). It has been hypothesized that such traits, including sperm storage, evolved independently several times, possibly as a more efficient investment of resources in deep sea environments, where locating partners can be more difficult (Quinn Jr. 1983). The independent position of Seguenzioidea in our topology is in agreement with this hypothesis.

Haliotoidea is the only superfamily currently assigned to Lepetellida (Bouchet et al. 2017; WoRMS 2019) that actually is not recovered in this vetigastropod order in our results. Instead, our phylogeny puts Haliotidae as the sister group to a clade of Lepetellida *s.s.* and Trochida (Figure 1.2). Even if one considers the alternative topology of the coalescent-based and partitioned ML analyses (Figures A.3, A.4), Haliotidae is still not included with the remaining Lepetellida, instead being sister to Trochida alone. Haliotoidea therefore appears to be an independent lineage of Vetigastropoda, not pertaining to any of the four currently recognized orders. The exact position of Haliotidae is however still not certain. Some previous studies based on morphological and a few molecular markers have found Haliotidae as closely related to Trochoidea (Aktipis and Giribet 2010; Geiger and Thacker 2005; Kano 2008; Ponder and Lindberg 1997), but our analyses indicate this could be a biased result (see subsection 1.4.1). Although Haliotidae does not present a particularly long branch in our tree, increasing taxon sampling for the family is a possible strategy to help determine its phylogenetic placement. Haliotidae has only one extant genus, which is why only one species was sampled here. Their fossil record is relatively recent, with most records being from the Cenozoic and only a few from the late Mesozoic (Groves and Alderson 2008). If the extant diversity of abalones is indeed a recent radiation, it is not clear how much further sampling could help in resolving the position of the family among vetigastropods.

Some previous molecular studies find Fissurelloidea as one of the lineages that first diverges from remaining vetigastropods (Aktipis and Giribet 2010; Kano 2008; Lee et al. 2016; Uribe et al. 2016). This is particularly worrisome in mitogenome analyses, where Fissurellidae always presents the longest branch in the tree, having a different gene arrangement and faster rates of mitochondrial evolution compared to other vetigastropods (Lee et al. 2016; Uribe et al. 2016, 2017). Such results indicate a likely effect of LBA pushing fissurellids closer to the outgroups. Our analyses, on the other hand, have an extensive sample of fissurellid taxa and support the placement of the family in Lepetellida *s.s.* as the sister group to a clade of Lepetodriloidea, Lepetelloidea and Scissurelloidea (Figure 1.2), a relationship that has been recovered before (Williams et al. 2008; Williams and Ozawa 2006). Within Fissurellidae, relationships are well resolved and congruent with the ones recently recovered in a study with denser taxon sampling, but fewer molecular markers (Cunha et al. 2019). Zeidorinae is sister group to the other sampled fissurellids, and Diodorinae and Fissurellinae form a clade. Emarginulinae is not monophyletic, with *Emarginula* being more closely related to Diodorinae and Fissurellinae than to other emarginuline.

Trochoidea is the most diverse superfamily of Vetigastropoda, with over 2,000 described species, and is here recovered as monophyletic with an extensive representation of ten out of 13 families. Early molecular results had elevated Phasianellidae and Angariidae to superfamilies Phasianelloidea and Angarioidea (Williams et al. 2008), finding that they represented earlier splits in the vetigastropod tree (Aktipis and Giribet 2011; Williams et al. 2008). Our results do not support this hypothesis, and instead agree with recent mitogenomic analyses that reinstated these families as members of Trochoidea (Uribe et al. 2017).

### 1.4.3 Relationships within Trochoidea

Although mitogenome studies have suffered from poor taxon sampling and long branches for many vetigastropod superfamilies, they have provided more stable and relatively better-sampled trees for Trochoidea (Lee et al. 2016; Uribe et al. 2016, 2017). Where sampling overlaps, our results are in agreement with mitogenomic analyses: Calliostomatidae and Trochidae are sister groups, and together they are sister group to a clade of Angariidae and Phasianellidae, Tegulidae (not monophyletic) and Turbinidae (Figure 1.2). We further sampled additional families, with over 75% of the family diversity being represented. With this sampling we find some similarities, but many discordances with more densely sampled studies of Trochoidea that used a handful of nuclear and mitochondrial genes (Williams 2012; Williams et al. 2010).

Skeneidae is the sister group to all other trochoids, as previously reported (Williams 2012). While past work has found Liotiidae closely related to Calliostomatide or Tegulidae (Aktipis and Giribet 2011; Kano 2008; Williams 2012; Williams et al. 2008), we recover Liotiidae as sister group to Angariidae (Figure 1.2). A similar clade had already been recovered before (Aktipis and Giribet 2011), but without support from other studies on trochoids.

Relationships of subfamilies within Trochidae were well resolved and are fully concordant with the latest and more densely sampled phylogeny based on a handful of markers (Williams 2012), providing additional support for the backbone tree of this diverse family. We also find that *Stomatella* is more closely related to *Stomatia* than to *Stomatolina*, as recovered in Williams et al. 2010.

Finally, Tegulidae is divided in two clades, one of which is more closely related to Turbinidae (Figure 1.2). This result has been found in other studies (Uribe et al. 2017; Williams 2012), but the clade including *Tectus* remains treated as unassigned, and a new family designation is necessary. Of the nine genera still classified as Tegulidae, six have been sampled here and in other recent studies, with *Cittarium, Rochia* and *Tectus* consistently recovered in the unassigned clade (Uribe et al. 2017; Williams 2012).

## 1.5 Conclusion

With an extensive sample of all vetigastropod superfamilies and about half of the modern familial diversity, we provide the first phylogenomic framework for deep relationships in Vetigastropoda. Our sampling also allows for a robust inference of backbone relationships within the most diverse vetigastropod clades: superfamily Trochoidea, and families Trochidae and Fissurellidae. Divergences at all levels are well supported and largely concordant between inference methods.

We find that Trochoidea (sole member of the order Trochida) is the sister group to Lepetellida *s.s.* (including Lepetodriloidea, Lepetelloidea, Scissurelloidea and Fissurelloidea). Haliotoidea is sister group to this large clade, which in turn is the sister group to Seguenzioidea. The position of Haliotoidea remains the most uncertain node in the tree. Additional sampling of abalones could potentially provide more confidence in the results recovered here, but because modern Haliotidae are a relatively recent radiation, it is unclear how much phylogenetic signal further sampling will add to these ancient divergences.

With the exception of the position of Pleurotomarioidea as the sister group to all other vetigastropods, our results differ considerably from topologies previously inferred for vetigastropods. In particular, the large taxon and gene sampling and the use of complex models of site-heterogeneity allowed us to resolve relationships that have been greatly affected by long branch attraction artifacts in previous studies. Such artifacts have been widespread in vetigastropod analyses, from datasets of a handful of molecular markers to complete mitochondrial genomes. Difficulty in resolving deep relationships in this group are likely due to the ancient and fast divergence between the main lineages, and the lack of power in previous datasets derived from insufficient sequence data. Our data illustrate that it is possible to reconstruct these ancient splits with a comprehensive sampling of taxa and sequence data.

## 1.6 Author’s contributions

TJC and GG conceived the study, collected and identified specimens. TJC carried out lab work, analyzed the data and drafted the manuscript. Both authors improved the manuscript and gave final approval for publication.

## 1.8 Acknowledgments

We are grateful to James Reimer for kindly providing support for fieldwork in Okinawa. We also thank Vanessa Knutson, Shawn Miller, Hin Boo Wee, Kristen Soong and Taku Ohara for help in the field. The computations in this paper were done on the Odyssey cluster supported by the FAS Division of Science, Research Computing Group at Harvard University.

## Appendix A

**FIGURE A.1:**
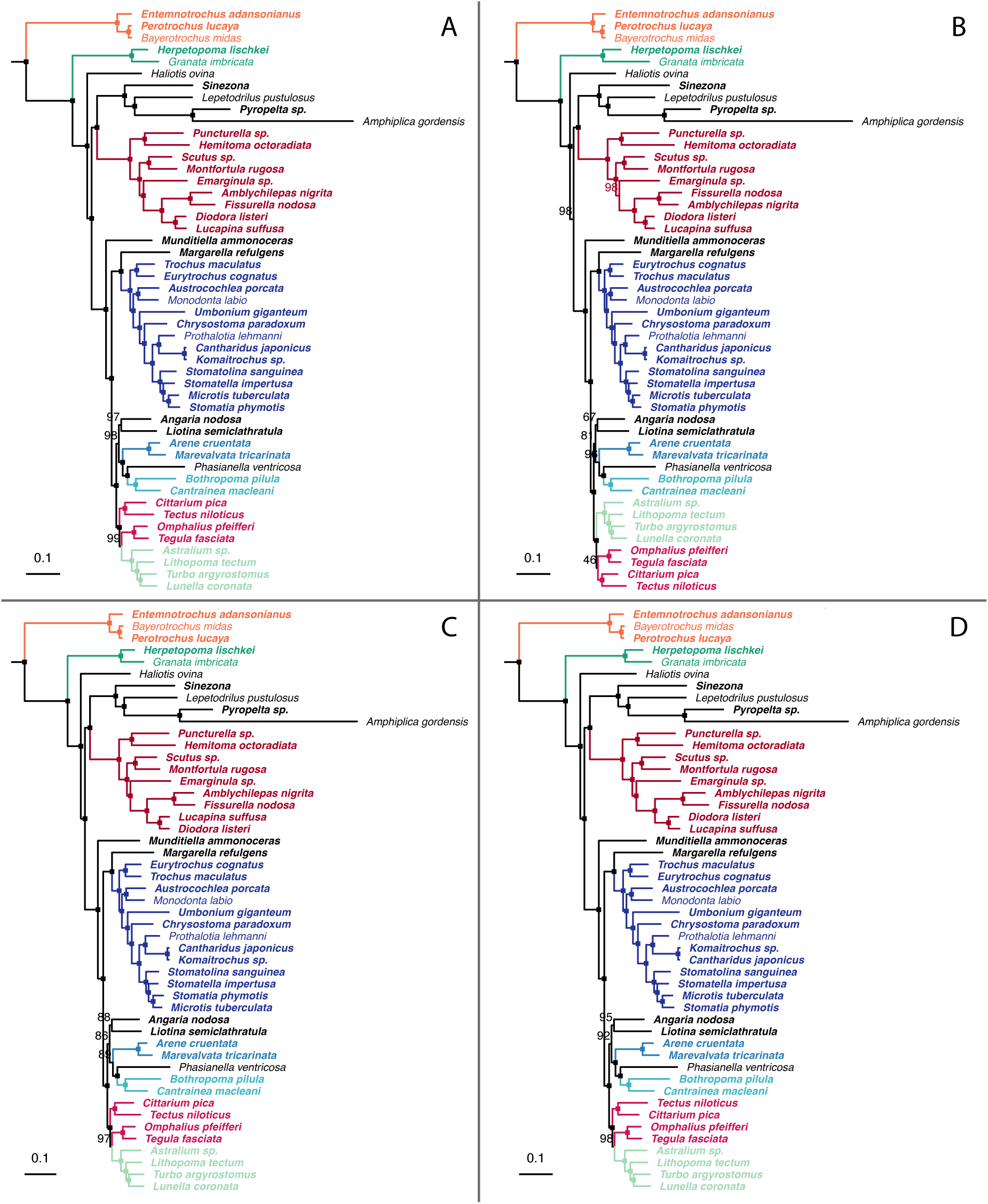
Vetigastropod phylogeny inferred with maximum likelihood and a profile mixture model on each of four matrices: (A) matrix 1, (B) matrix 2, (C) matrix 3, (D) matrix 4. Single squares mark branches with full support; remaining nodes show the bootstrap value. New transcriptomes in bold. See Methods for details.

**FIGURE A.2:**
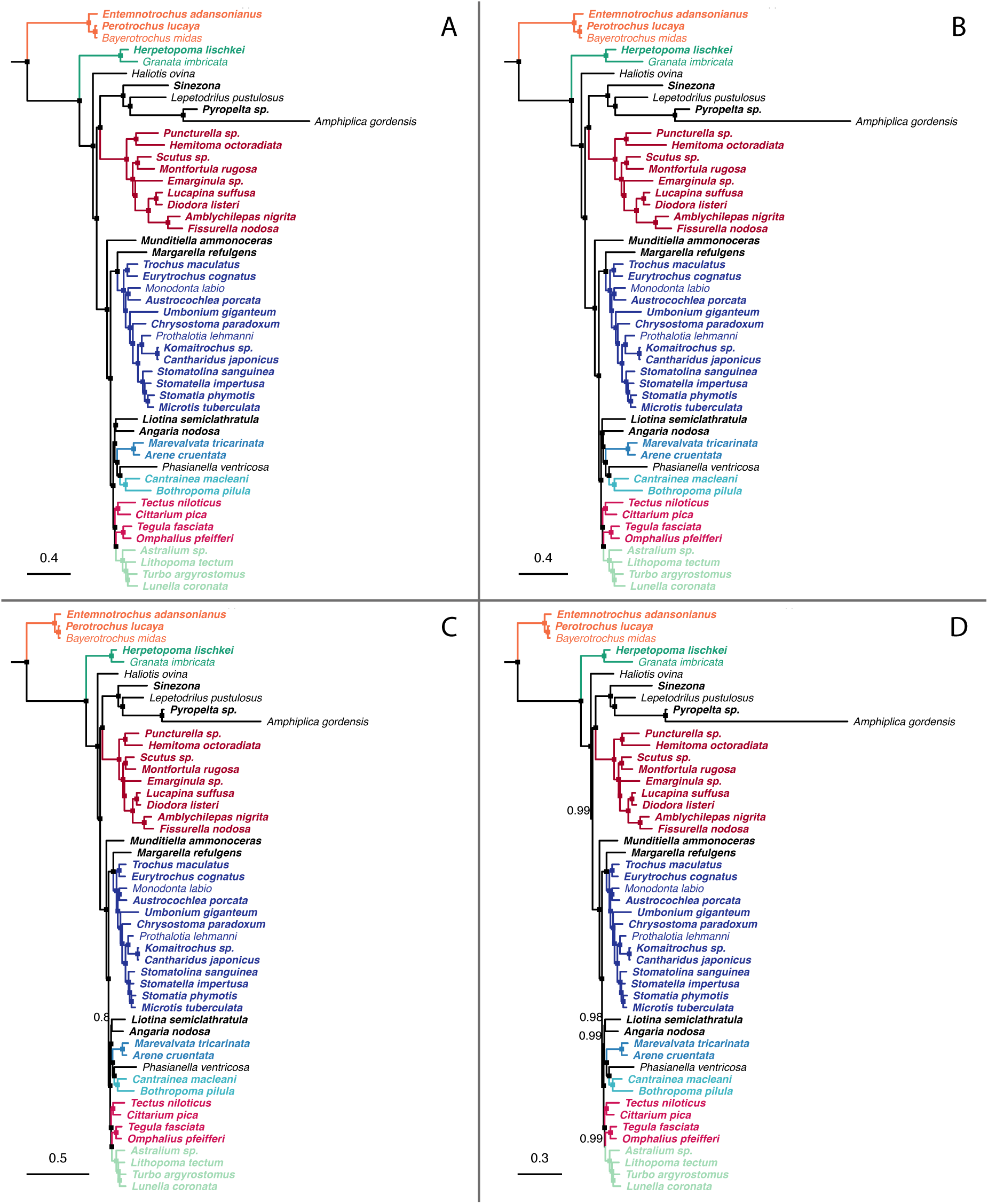
Vetigastropod phylogeny inferred with Bayesian inference on each of four matrices: (A) matrix 1, (B) matrix 2, (C) matrix 3, (D) matrix 4. Single squares mark branches with full support; remaining nodes show the posterior probability. New transcriptomes in bold. See Methods for details.

**FIGURE A.3:**
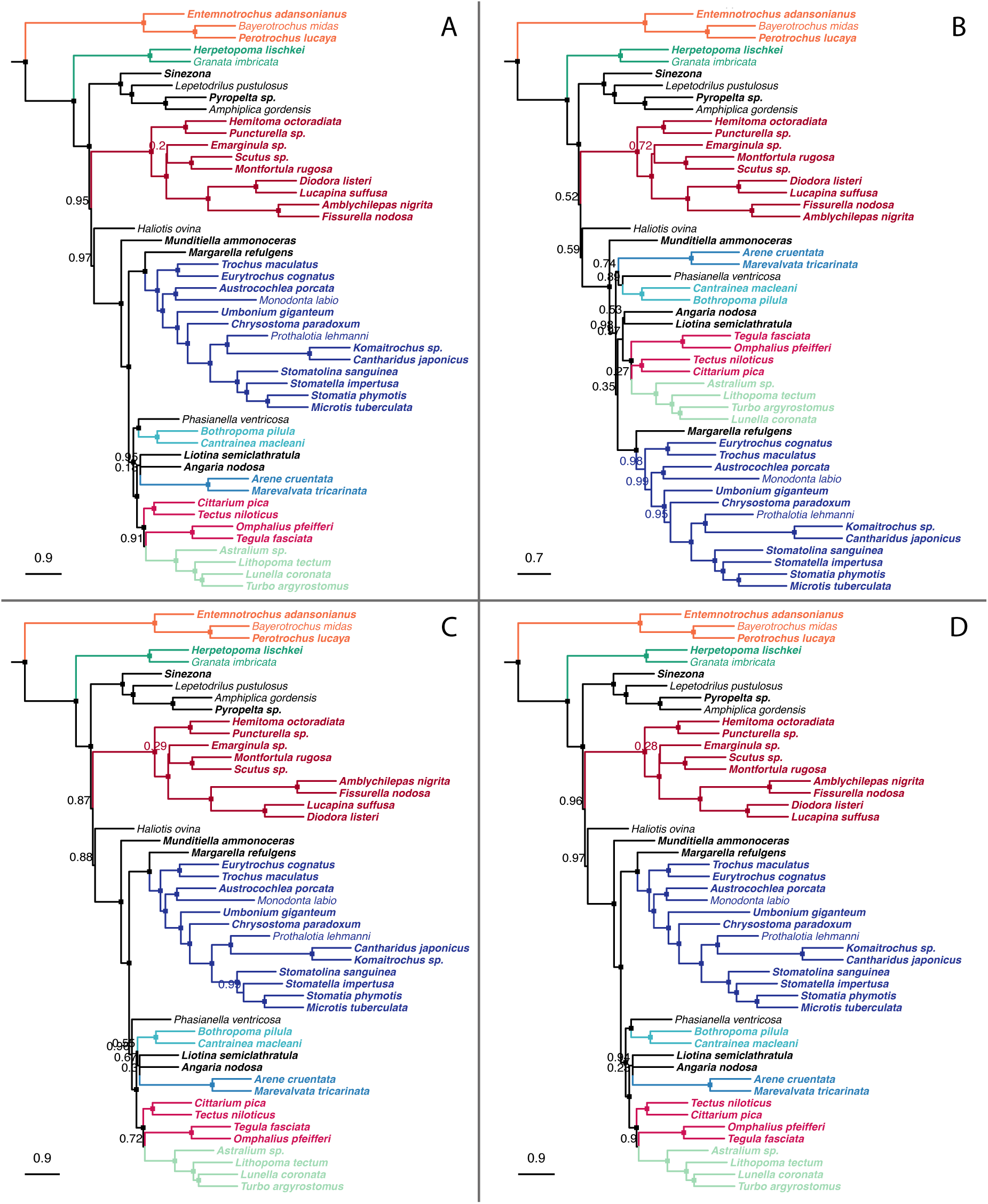
Vetigastropod phylogeny inferred with a shortcut coalescent-based method on each of four matrices: (A) matrix 1, (B) matrix 2, (C) matrix 3, (D) matrix 4. Single squares mark branches with full support; remaining nodes show the branch support value. New transcriptomes in bold. See Methods for details.

**FIGURE A.4:**
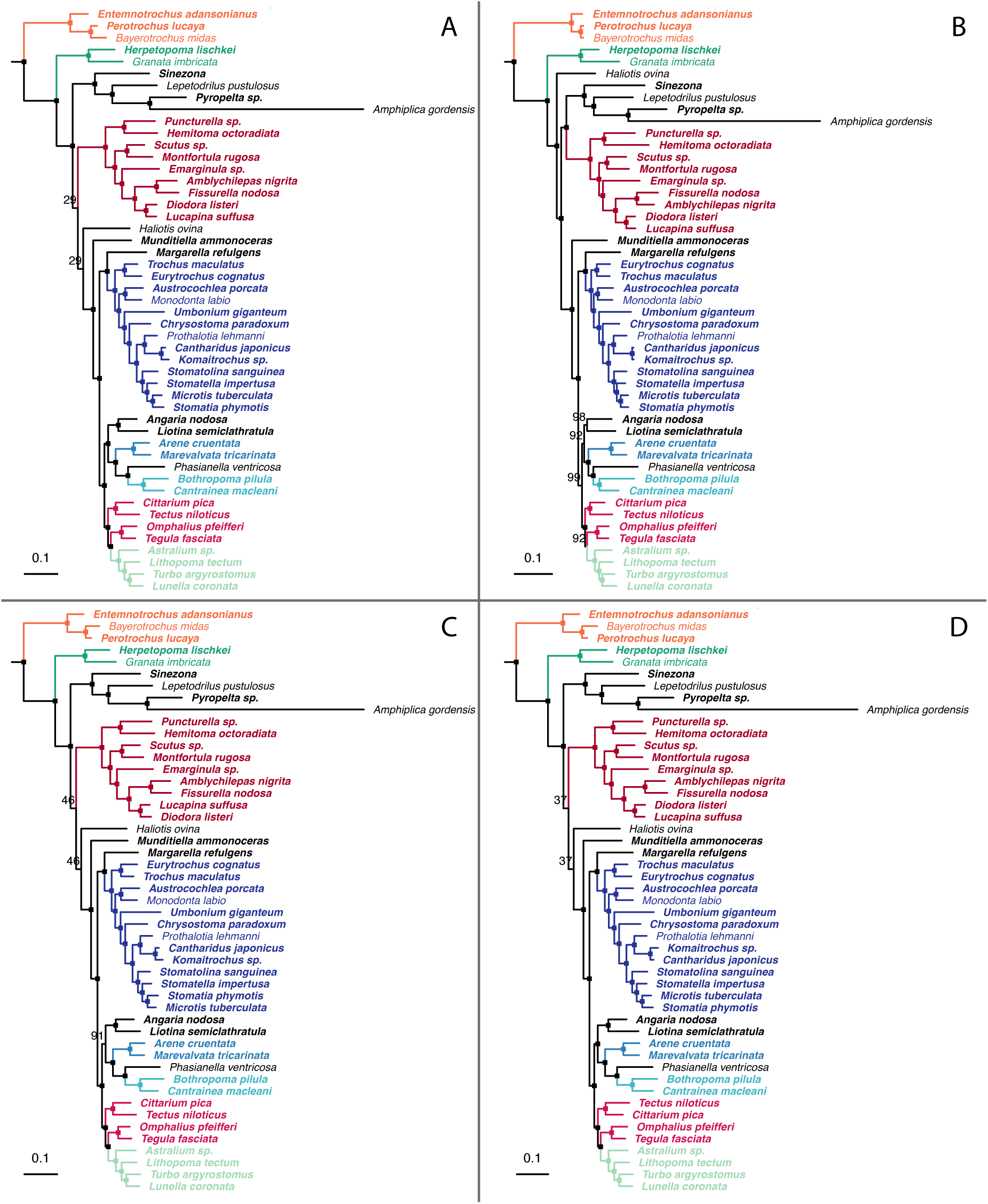
Vetigastropod phylogeny inferred with maximum likelihood and genewise partitioning on each of four matrices: (A) matrix 1, (B) matrix 2, (C) matrix 3, (D) matrix 4. Single squares mark branches with full support; remaining nodes show the bootstrap value. New transcriptomes in bold. See Methods for details.

